# Memory Image Completion (MIC): Establishing a task to behaviorally assess pattern completion in humans

**DOI:** 10.1101/366203

**Authors:** Paula Vieweg, Martin Riemer, David Berron, Thomas Wolbers

## Abstract

For memory retrieval, pattern completion is a crucial process that restores memories from partial or degraded cues. Neurocognitive aging models suggest that the aged memory system is biased toward pattern completion, resulting in a behavioral preference for retrieval over encoding of memories. While there are behavioral tasks to assess the encoding side of these memory differences, pattern completion has received less attention in the literature. Here, we built on our previously developed behavioral recognition memory paradigm – the Memory Image Completion task (MIC) – a task to specifically target pattern completion. First, we used the original design with concurrent eye-tracking in order to rule out perceptual confounds that could interact with recognition performance. Second, we developed parallel versions of the task to accommodate test settings in clinical environments or longitudinal studies. The results show that older adults have a deficit in pattern completion ability with a concurrent bias toward pattern completion – a replication of previously found effects. Importantly, eye-tracking data during encoding could not account for age-related performance differences. At retrieval, spatial viewing patterns for both age groups were more driven by stimulus identity than by response choice, but compared to young adults, older adults’ fixation patterns overlapped more between stimuli that they (wrongly) thought had the same identity. This supports the observation that when making errors older adults choose responses perceived as similar to the correct stimulus, which is interpreted as a bias toward pattern completion. Additionally, two shorter versions of the task yielded comparable results, and no general learning effects were observed for repeated testing. Together, we present evidence that the MIC is a reliable behavioral task that targets pattern completion, that is easily and repeatedly applicable, and that is made freely available online.

## Introduction

In everyday life, we constantly recognize people, places and objects. Often, the cues that trigger such retrieval are incomplete or different from their original encounter. For example, imagine walking down an avenue of poplars during winter, when you first experienced them blossoming in spring. Despite the lack of leaves and blossoms, recognition of this as the same avenue is seamlessly enabled by a process called pattern completion.

Pattern completion and separation have received widespread attention in neuroscience to foster the understanding of how the brain encodes and retrieves memories (for extensive reviews see Yassa and Stark, 2011; Hunsaker and Kesner, 2013). Based on computational models (Marr, 1971; McClelland, 1994; Treves and Rolls, 1994; O’Reilly et al., 1998; Hasselmo and McClelland, 1999) and findings in rodent studies (Nakazawa et al., 2002; Guzowski et al., 2004; Leutgeb et al., 2004, 2007; Vazdarjanova and Guzowski, 2004; Leutgeb and Leutgeb, 2007; Neunuebel and Knierim, 2014), pattern completion is thought to restore memory traces from partial or degraded input via neural reactivation, mainly originating in hippocampal region CA3. In contrast, pattern separation is considered as the complimentary process of differentiating very similar memories via orthogonalization of neural representations, which predominantly occurs in the dentate gyrus (DG).

While both processes are defined based on neural computations, their behavioral outcomes have been approximated in human research with behavioral tasks. A central paradigm is the Mnemonic Similarity Task (MST; Stark et al., 2015), in which participants are presented with very similar stimuli (lures) amongst novel and repeated stimuli (Kirwan and Stark, 2007; Bakker et al., 2008; Lacy et al., 2011; etc.). While the MST efficiently induces memory interference, the use of highly similar stimuli during encoding selectively favors pattern separation because similar memory traces need to be differentiated. It is possible that pattern completion is merely a secondary finding in these studies, i.e., without being explicitly manipulated. Hunsaker and Kesner (2013 p. 40) have indeed suggested that the presentation of parts of an original cue may engage pattern completion more independently than the presentation of similar cues. Additionally, they suggested that even though both processes likely contribute to memory encoding and retrieval, pattern separation may be more involved in the former and pattern completion in the latter. Thus, in order to increase the impact of either pattern separation or completion, studies should focus on the respective corresponding phase of memory processing. This idea received additional support from an eye-tracking study (Molitor et al., 2014), showing that false alarm trials in the MST (i.e., a similar stimulus judged as old), which were interpreted to involve pattern completion, were associated with fewer fixations during encoding. This observation suggests an additional involvement of pattern separation processes.

Accommodating the above suggestions, we have previously developed the Memory Image Completion task (MIC; Vieweg et al., 2015; named MIC in Baker et al., 2016). In this paradigm, gradually less complete versions of a learned visual stimulus reduce recognition performance, which was interpreted to reflect an increase in pattern completion demands. The practical suitability of our task was later independently confirmed in a comparative meta analysis (Liu et al., 2016), and it has also been employed successfully in a lesion study (Baker et al., 2016). A second critical aim of the MIC was the assessment of distinct age effects. The model of neurocognitive aging by Wilson et al. (2006) suggests that the aged memory system should show a bias toward pattern completion and a concurrent deficit in pattern separation. Consistent with this prediction, pattern separation has been shown to be less efficient in aging (Toner et al., 2009), accompanied by CA3/DG hyperactivity and rigidity (Yassa et al., 2011a; b), and it is more sensitive to cognitive decline than standard recognition memory (Holden et al., 2013; Stark et al., 2013). Complementary, our previous results suggested deficient but reinforced pattern completion (Vieweg et al., 2015).

In addition to behavioral performance, viewing behavior can inform about memory-related processing. For example, higher fixation numbers during encoding have been suggested to reflect increased accumulation of information, leading to better memory representations (Pertzov et al., 2009) and hence increased memory performance (Loftus, 1972). This can be important when interpreting behavioral results in pattern separation/completion tasks, which have attributed age-related performance changes to a shift from pattern separation to pattern completion during memory retrieval (Yassa et al., 2011b). However, Molitor et al. (2014) found fewer fixations on falsely recognized lure items during encoding, suggesting that a purely retrieval-based process is unlikely or at least insufficient to explain an observed behavioral bias.

To exhaust the possibilities of a task specifically tailored to pattern completion processes, we sought to refine and expand the MIC. First of all, we wanted to replicate the previous findings and rule out contributions of perceptual confounds. Even though the paradigm does not use incidental encoding and accommodates potential encoding differences through a learning criterion, we investigated whether specific viewing patterns could explain any age-related retrieval differences. This question was addressed with an eye-tracking study that characterized eye movements associated with the MIC and their relation to cognitive aging. Secondly, for the MIC to be useful in longitudinal studies and for clinical research, it needs to be short, and parallel versions should be available. Therefore, in a second experiment we developed four shorter versions of the MIC and assessed performance stability over multiple testing occasions.

## Experiment 1

### Methods

The task administered here parallels the Memory Image Completion task (MIC) we have used in our previous work and which is described in more detail there (Vieweg et al., 2015; Baker et al., 2016). Concurrently, we used eye-tracking to monitor participants’ viewing behavior during the task.

#### Subjects

All participants were recruited from existing databases at the German Center for Neurodegenerative Diseases (DZNE) in Magdeburg. They underwent several neuropsychological tests and health assessments prior to the experiment: health questionnaire (Diersch, 2013), Montreal Cognitive Assessment (MoCA; Nasreddine et al., 2005), multiple choice vocabulary test (MWT-B; Lehrl et al., 1995), digit symbol substitution test (DSST; Wechsler, 2008), Rey-Osterrieth Complex Figure (ROCF) Test as copy and 30 minutes delayed recall (DR; Rey, 1941; Corwin and Bylsma, 1993); their visual dominance was assessed, as well as subjective eyesight (using German grading system 1 “very good” – 6 “insufficient”) and objective eyesight (visual acuity determined on a pocket card test). Participants who scored less than 23 on the MoCA (N = 2; Luis et al., 2009), who had chronic psychological or neurological problems (N = 1), who had insufficient eyesight (N = 3), or whose eyes could not be tracked due to their glasses (N = 4; e.g. reflection, small frame, dark frame), were excluded from the study. The remaining 26 young (21-35 years; 13 females; 6 wore glasses, 7 wore contact lenses) and 24 older adults (63-77 years; 12 females, 22 wore glasses) were healthy and had neuropsychological scores characteristic of their age group, including increased vocabulary scores and reduced delayed recall performance (see Table 1). These findings are common in aging studies (Lövdén et al., 2005) and authenticate the cognitive typicality of our sample.

**Table 1.**
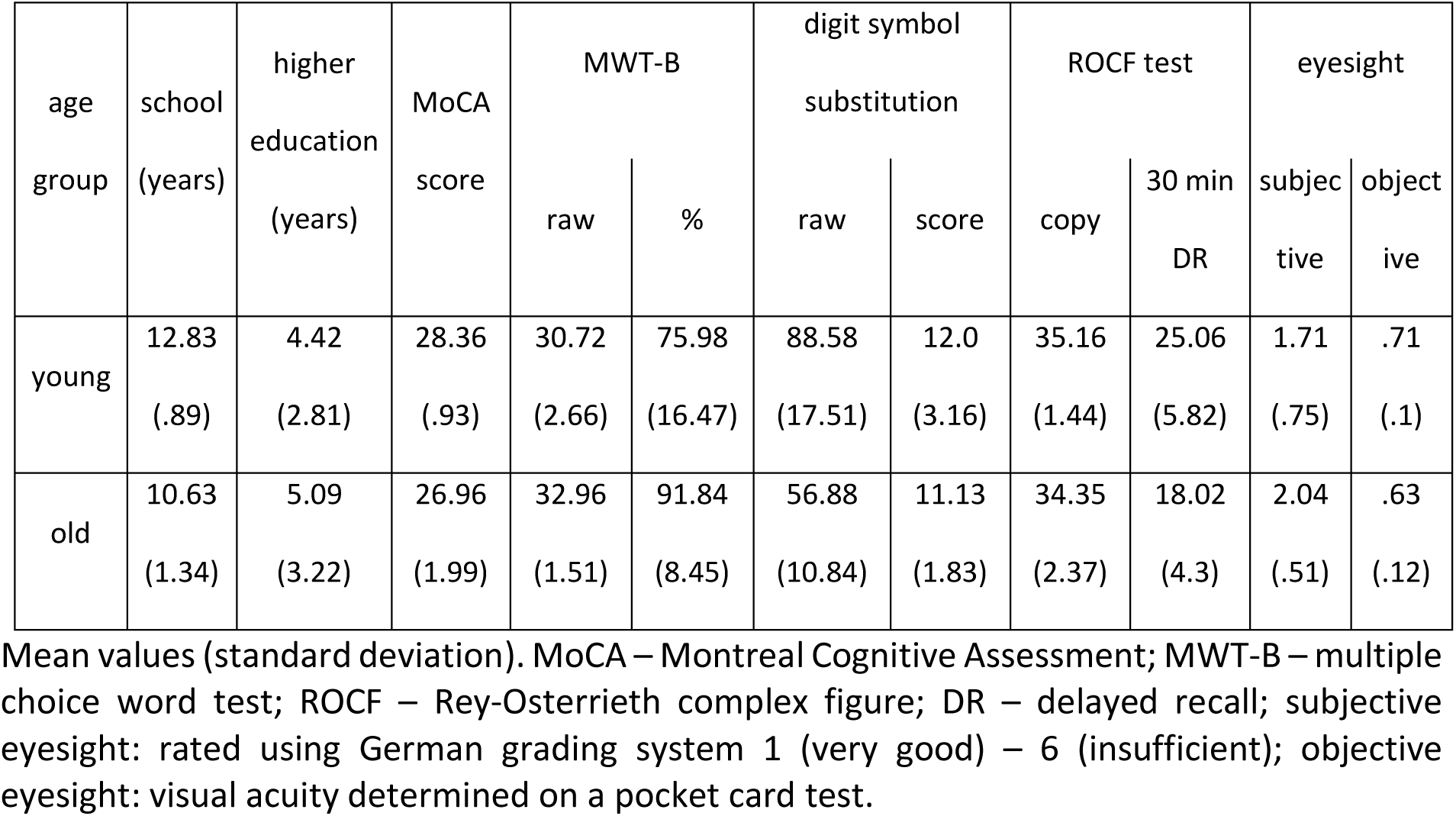
Experiment 1 - Health questionnaire and neuropsychological data.

| age group | school (years) | higher education (years) | MoCA score | MWT-B |  | digit symbol substitution |  | ROCF test |  | eyesight |  |
| --- | --- | --- | --- | --- | --- | --- | --- | --- | --- | --- | --- |
|  |  |  |  | raw | % | raw | score | copy | 30 min DR | subjective | objective |
| young | 12.83<br>(.89) | 4.42<br>(2.81) | 28.36<br>(.93) | 30.72<br>(2.66) | 75.98<br>(16.47) | 88.58<br>(17.51) | 12.0<br>(3.16) | 35.16<br>(1.44) | 25.06<br>(5.82) | 1.71<br>(.75) | .71<br>(.1) |
| old | 10.63<br>(1.34) | 5.09<br>(3.22) | 26.96<br>(1.99) | 32.96<br>(1.51) | 91.84<br>(8.45) | 56.88<br>(10.84) | 11.13<br>(1.83) | 34.35<br>(2.37) | 18.02<br>(4.3) | 2.04<br>(.51) | .63<br>(.12) |

Mean values (standard deviation). MoCA – Montreal Cognitive Assessment; MWT-B – multiple choice word test; ROCF – Rey-Osterrieth complex figure; DR – delayed recall; subjective eyesight: rated using German grading system 1 (very good) – 6 (insufficient); objective eyesight: visual acuity determined on a pocket card test.

Informed written consent was obtained before the experiment, and the study received approval from the Ethics Committee of the University of Magdeburg. All participants received monetary compensation of 6.50€/h.

#### Materials

We used 15 black and white line-drawn images of simple indoor scenes from Hollingworth and Henderson (1998; 5 to be learned, 5 to test learning, 5 new). Stimulus completeness was manipulated by gradually masking all stimuli used in the test phase with a grid of white circles resulting in five different completeness levels (100%, 35%, 21%, 12%, and 5%; percentages reflect the amount of the image visible through the mask; see Figure 1A). Including the complete stimuli, this resulted in a total of 50 stimuli in the test phase (10 images × 5 completeness levels), each of which was repeated four times.

**Figure 1.**
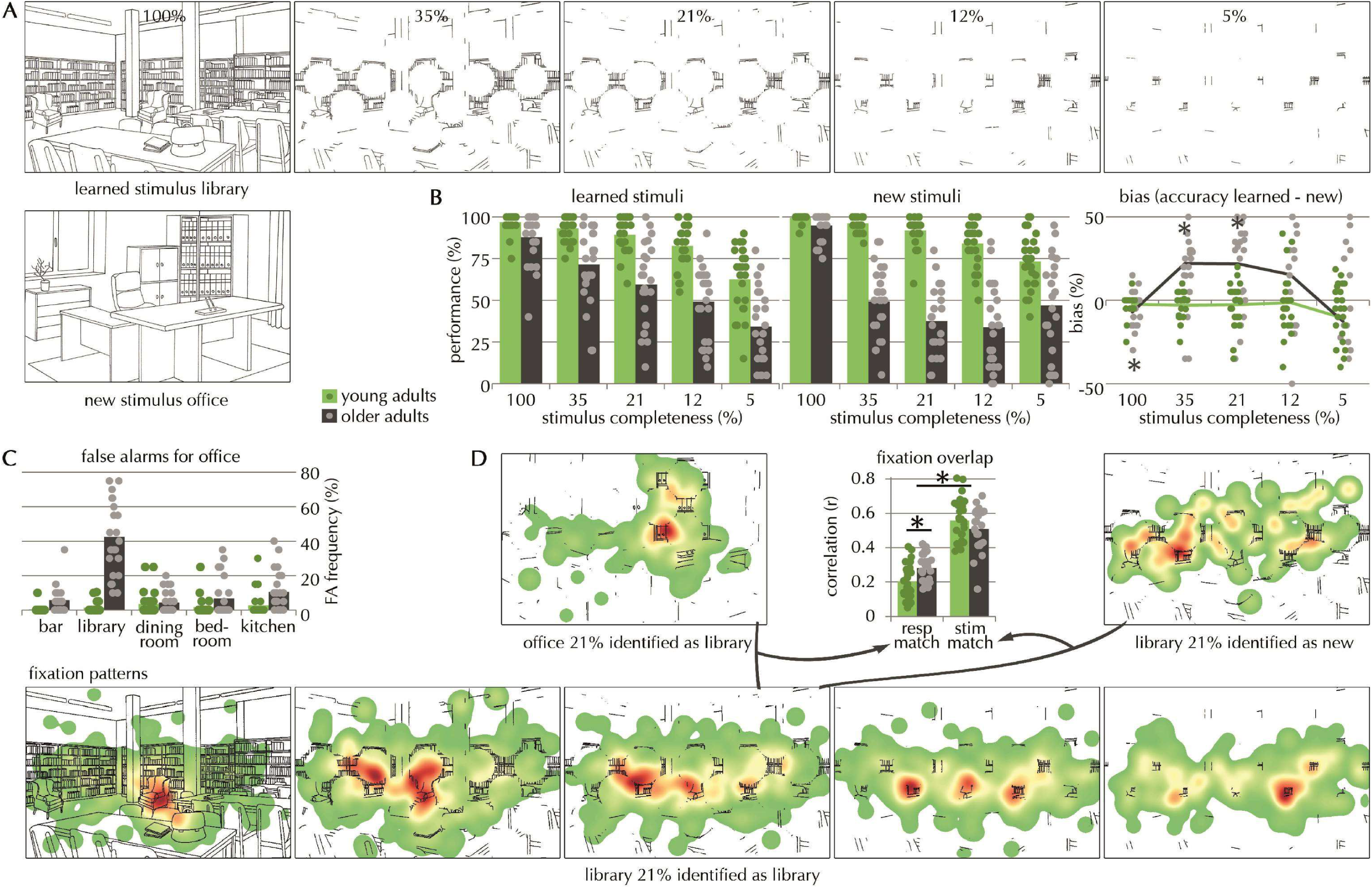
- Experiment 1. **(A)** learned stimulus library in all possible levels of completeness (percentages indicate portions of the image that are still visible through the mask) and new stimulus office. **(B)** Left: performance for both age groups (green: young adults, gray: older adults), separately for learned and new stimuli for the 5 different levels of stimulus completeness (mean); right: bias measure (see methods for a detailed explanation) - difference in accuracy scores for learned minus new stimuli calculated separately for each participant (mean); positive values indicate a bias toward pattern completion, significant differences from 0 are indicated with * separately for each age group as indicated by color. **(C)** exemplary false alarm distribution for new stimulus office showing that older adults chose one particular false response option (library) most often rather than guessed more overall, which would have led to similar frequencies for all 5 response options. **(D)** exemplary fixation heatmaps, warmer colors indicate higher fixation numbers; lower panel: fixation heatmaps for correctly identified stimuli (exemplary: library identified as library) for each masking level (100 – 5%); top left: fixation heatmap for a response-matching erroneous response (exemplary: office 21% identified as library), top right: fixation heatmap for a stimulus-matching erroneous response (exemplary: library 21% identified as kitchen); top middle: fixation overlap - fixation heatmaps for correctly identified stimuli were correlated with heatmaps for stimulus-matching erroneous responses (stim match), or with response-matching erroneous responses (resp match). Pearson’s correlation coefficients were Fisher’s z-transformed and averaged across stimuli and completeness levels, separately for both age groups (correlation values in the figure were back-transformed for better comprehensibility), significant differences are indicated with *: stim-match correlations were overall higher than resp-match correlations suggesting that the viewing patterns were driven by the stimulus rather than by the response choice, but older adults’ fixation patterns for resp-match stimuli correlated more than those of young adults (and these fixation overlap scores correlated with the response bias in older adults; see results for details) suggesting that the more likely a participant is to falsely recognize the office as the library, the more the fixation patterns for library and office overlap.

#### Procedure

In a short study phase, participants had to learn a set of five different line-drawn scenes. Each item was presented randomly with a preceding semantic stimulus label. After learning, we verified that participants had learned the items by testing whether they could correctly identify them among new items until reaching a learning criterion (three consecutive correct identifications). In the main test phase of the experiment, participants had to consecutively judge the identity of a given stimulus out of a fixed set of response options (all of the learned stimuli’s labels and ‘none of these’; chance level: 1/6). A presented stimulus could either be a learned or a new stimulus, and was either presented in full (100%) or in a gradually masked version (35%, 21%, 12%, and 5%). Each stimulus was randomly shown four times and for two seconds. Responses were self-paced, and participants were asked to respond accurately rather than quickly. Responses were scored as correct only when participants had picked the correct name for a learned stimulus or ‘none of these’ for a new stimulus.

Additionally, eye movements were recorded during image presentation in all parts of the experiment. At the beginning of each phase of the experiment, gaze position was calibrated with a 9-fold grid of fixation points. Participants were asked to refrain from looking down at the keyboard when responding, because head movements could cause slight shifts of the eye-tracker. However, as participants had to perform several button presses with at least five response options per trial, this could not be prevented entirely. Therefore, every trial started with a drift correction prior to image presentation, in which participants had to fixate a small white circle in the middle of the screen. If drift correction exceeded a visual angle of 5°, tracking of gaze position was adjusted by recalibration. During the test phase, gaze position was recalibrated every 70 trials by default. Gaze position was always recorded from the dominant eye (32 right, 18 left), except for two participants, for whom the recorded eye was switched after half of the experiment due to recalibration failure. Eye-movement data of two young participants were excluded for the first half of the study phase due to a calibration error.

#### Eye-tracking acquisition and analyses

Eye movements were recorded with a head-mounted EyeLink II tracker (SR Research, Ontario, Canada) at a sampling rate of 500 Hertz. Participants were seated 50 centimeters in front of a 15 inch computer screen (1024 × 768 pixel resolution, 60 Hertz refresh rate). The experiment was programmed in Matlab 2013a with the Psychtoolbox Add-on to integrate the eye-tracker. Eye movements were recorded for each participant, and saccade, blink, and fixation data were calculated as follows. Blinks were defined as missing pupil data over three consecutive samples. Fixations were defined as static gaze positions, i.e., no movements greater than 0.1° visual angle, and faster than 30°/s velocity and 8000°/s^2^ acceleration. Saccades consisted of all other recordings. Eye-tracking analyses focused on number and duration of fixations during the two second image presentation at study and test, and their proportion in pre-defined interest areas (IAs). IAs were equivalent to the inverse masking grid used to manipulate stimulus completeness, i.e., they consisted of all the areas in the image that were still visible through the mask, including an extended margin of 25 pixels to include areas that were in the foveal focus (˜1.5° visual angle).

To investigate the spatial distribution of fixations, fixation heatmaps were extracted per trial for each participant, i.e., a matrix containing fixation numbers at each image pixel coordinate. Each fixation’s foveal area was also included into the fixation matrix; it was defined as ~1.5° visual angle around the fixation (25 pixel radius). Heatmaps were then averaged across repetitions and smoothed with a Gaussian filter (50 × 50 pixels), resulting in one fixation heatmap per condition (accounting for stimulus identity, completeness and response made, see Figure 1D). Two-dimensional Pearson’s correlation coefficients were then calculated between condition-specific heatmaps.

#### Exploration of group differences using Bayesian analyses

Complementary to standard frequentist statistics (t-tests, analyses of variance - ANOVA), we performed Bayesian analyses to estimate support for the null hypotheses, because the absence of a significant difference between two measures in classical frequentist statistics cannot directly be taken to indicate that the two measures are in fact not different. Consequently, we calculated Bayes factors in JASP Version 0.8.4.0 (jasp-stats.org) using Bayesian t-tests and ANOVAs with default settings (t-test: Cauchy prior width 0.707; ANOVA: prior fixed effects 0.5, random effects 1, covariates 0.354) providing an indication of the strength of evidence in favor of the null hypothesis, i.e., if there were no differences between age groups. The Bayes factor comparing the null hypothesis to the alternative hypothesis (*BF_01_*) means that the data are BF times more likely under the null than the alternative hypothesis. *BF_01_* values much greater than 1 support the conclusion that there is strong evidence in favor of the null hypothesis. We report *BF_01_* only (and not *BF_10_*) because our main focus was on the likelihood of the groups being similar. Note, though, that the null hypothesis is inherently more difficult to prove than the alternative, i.e. with increasing observations the BF converges towards the true hypothesis (null or alternative), but to reach moderate (BF > 3) or strong (BF > 10) evidence, considerably more data are required than for the alternative (see minimal BF in Felix Schönbrodt’s blog entry: http://www.nicebread.de/what-does-a-bayes-factor-feel-like/).

## Results

### Behavioral results

#### Accuracy

The following results were obtained by a three-way mixed ANOVA with factors age (young/old), stimulus completeness (100%, 35%, 21%, 12%, 5%), and stimulus type (learned/new). Young participants performed better than older participants (main effect of age: *F*_(1,48)_ = 142.01 *p* < .001). Reduced stimulus completeness resulted in less accurate performance (main effect of stimulus completeness: *F*_(4,192)_ = 153.102, *p* < .001), and this decrease was more pronounced in older adults (age × stimulus completeness: *F*_(4,192)_ = 33.582, *p* < .001). Performance per stimulus type did not differ overall (main effect of stimulus type: *F*_(1,48)_ = 0.568, *p* = .455), but interactions revealed that older participants were less accurate for new stimuli compared to learned stimuli (age × stimulus type: *F*_(1,48)_ = 5.408, *p* = .024; age × stimulus type × stimulus completeness: *F*_(4,192)_ = 14.582, *p* < .001). Post-hoc t-tests showed age group differences in performance across all levels of stimulus completeness for both learned and new stimuli (after Holm-Bonferroni multiple comparisons correction; all *p* < .01; level 100%: *t_learned_*_(48)_ = 2.887, *t*_new(48)_ = 2.936, level 35%: *t_learned_*_(48)_ = 4.667, *t_new_*_(48)_ = 11.045, level 21%: *t_learned_*_(48)_ = 5.676, *t_new_*_(48)_ = 12.418, level 12%: *t_learned_*_(48)_ = 6.508, *t_new_*(48) = 9.196, level 5%: *t_learned_*_(48)_ = 4.774, *t_new_*_(48)_ = 4.024). In summary, both age groups’ recognition ability decreased with reduced stimulus completeness, and older adults’ recognition ability was impaired in comparison to young adults, especially for new stimuli (see Figure 1B).

#### Response bias

We investigated potential response biases by subtracting individual accuracy scores for new stimuli from those for learned stimuli (see also Vieweg et al., 2015). Positive scores are indicative of better performance for to-be-retrieved (learned) stimuli suggesting a bias towards pattern completion, while negative scores indicate better performance for to-be-encoded (new) stimuli in turn suggesting a bias towards pattern separation.

A mixed ANOVA (age × stimulus completeness) revealed that stimulus completeness influenced the response bias (main effect of stimulus completeness: *F*_(4,192)_ = 14.582, *p* < .001), and older adults showed a more positive response bias than young adults (see Figure 1B, right; main effect of age: *F*_(1,48)_ = 5.408, *p* = .024). A two-way interaction indicated that stimulus completeness differentially affected the response bias dependent on the age group (age × stimulus completeness: *F*_(4,192)_ = 8.053, *p* < .001). Follow-up t-tests demonstrated between-group differences for the middle three completeness levels, but after Holm-Bonferroni multiple comparisons correction the 12% level did not reach significance (level 35%: *t*_(48)_ = − 4.68, *p* < .001, level 21%: *t*_(48)_ = −4.042, *p* < .001; level 12%: *t*_(48)_ = −2.107, *p* = .04).

We examined at which levels of stimulus completeness the scores established a response bias. To that end, per group, five one-sample t-tests of the average response bias scores against 0 revealed that older adults showed a positive response bias for completeness levels 35% and 21%, indicative of a pattern completion bias, and a negative response bias for level 100% after Holm-Bonferroni multiple comparisons corrections (see Figure 1B; level 100%: *t*_(23)_ = −2.416, *p* = .024; level 35%: *t*_(23)_ = 4.149, *p* < .001, level 21%: *t*_(23)_ = 3.819, *p* = .001; level 12%: *t*_(23)_ = 2.081, *p* = .049; level 5%: *t*_(23)_ = −1.392, *p* = .177). Young participants showed no response bias at all (all *p >* .05).

If participants completed towards the stimulus perceived as most similar, one false response option should have been chosen over all other false options. Alternatively, if participants were merely guessing, all false response options should occur equally often. Therefore, we investigated older adults’ distribution of false alarm options for new stimuli by calculating the group-average frequencies of all false choice options. Frequencies were then sorted from most (FA 1) to least (FA 5) chosen option for each new stimulus, and averaged across all new stimuli afterwards. A χ^2^-test of goodness-of-fit on the 5 false alarm options (*χ^2^*_(_*_4_*_)_ = 396.121, *p* < .001) revealed that participants indeed chose one particular false response option most often and were not randomly guessing (see Table 2). We inspected whether there was one dominant option for each stimulus, by contrasting the most frequent false alarm against the average of the other false alarms per stimulus (see Figure 1C for one example; stimulus ‘office’: *χ^2^*_(1)_ = 126.863, *p* < .001; stimulus ‘class room’: *χ^2^*_(1)_ = 19.862, *p* < .001; stimulus ‘restaurant’: *χ^2^*_(1)_ = 16.03, *p* < .001; stimulus ‘locker room’: *χ^2^*_(1)_ = 11.449, *p* = .001; stimulus ‘living room’: *χ^2^*_(1)_ = 2.462, *p* = .117). This indicates that older participants mostly completed towards the stimulus perceived as most similar.

**Table 2.** Experiment 1 - Response distribution for new stimuli.

| age group | CR | FA1 | FA2 | FA3 | FA4 | FA5 |
| --- | --- | --- | --- | --- | --- | --- |
| young | .89 (.07) | .02 (.02) | .04 (.04) | .03 (.03) | .01 (.02) | .01 (.01) |
| old | .43 (.2) | .26 (.14) | .13 (.07) | .09 (.05) | .06 (.05) | .05 (.03) |

Mean values (standard deviation). CR – correct rejection; FA – false alarm

#### Summary of the performance results

The present results replicated the findings from our previous study (Vieweg et al., 2015); both age groups’ recognition memory declined with decreasing stimulus completeness, older adults showed a stronger decline and they chose familiar responses over new ones, suggesting a bias towards pattern completion. Note that reaction times and confidence ratings followed the same profile as the original study (see Supplementary material).

### Eye-tracking results

#### Fixations during encoding

To investigate whether behavioral performance in the MIC could be driven by encoding effects, we examined fixations during the study phase. Although fixation durations did not differ between age groups (*t*_(48)_ = 0.715, *p* = .478, *BF_01_* = 2.867), younger adults fixated slightly more often on the stimuli than older adults (*t*_(48)_ = 2.053, *p* = .048, *BF_01_* = 0.650; see Table 3). In order to explore whether this difference was responsible for the age-related response bias during retrieval, we correlated the number of fixations during encoding with a cumulative partial response bias score per participant (mean of all partial response biases; as in Baker et al., 2016). In neither of the age groups did these measures correlate (Pearson correlation; young adults: *r* = −0.282, *p* = .163; older adults: *r* = −0.090, *p* = .674) suggesting that encoding differences were not predictive of response biases. Similarly, the older adults’ reduced recognition of learned items was not driven by differential encoding either (Pearson correlation with cumulative performance for partial learned items; young adults: *r* = 0.073, *p* = .721; older adults: *r* = 0.060, *p* = .758).

**Table 3.**
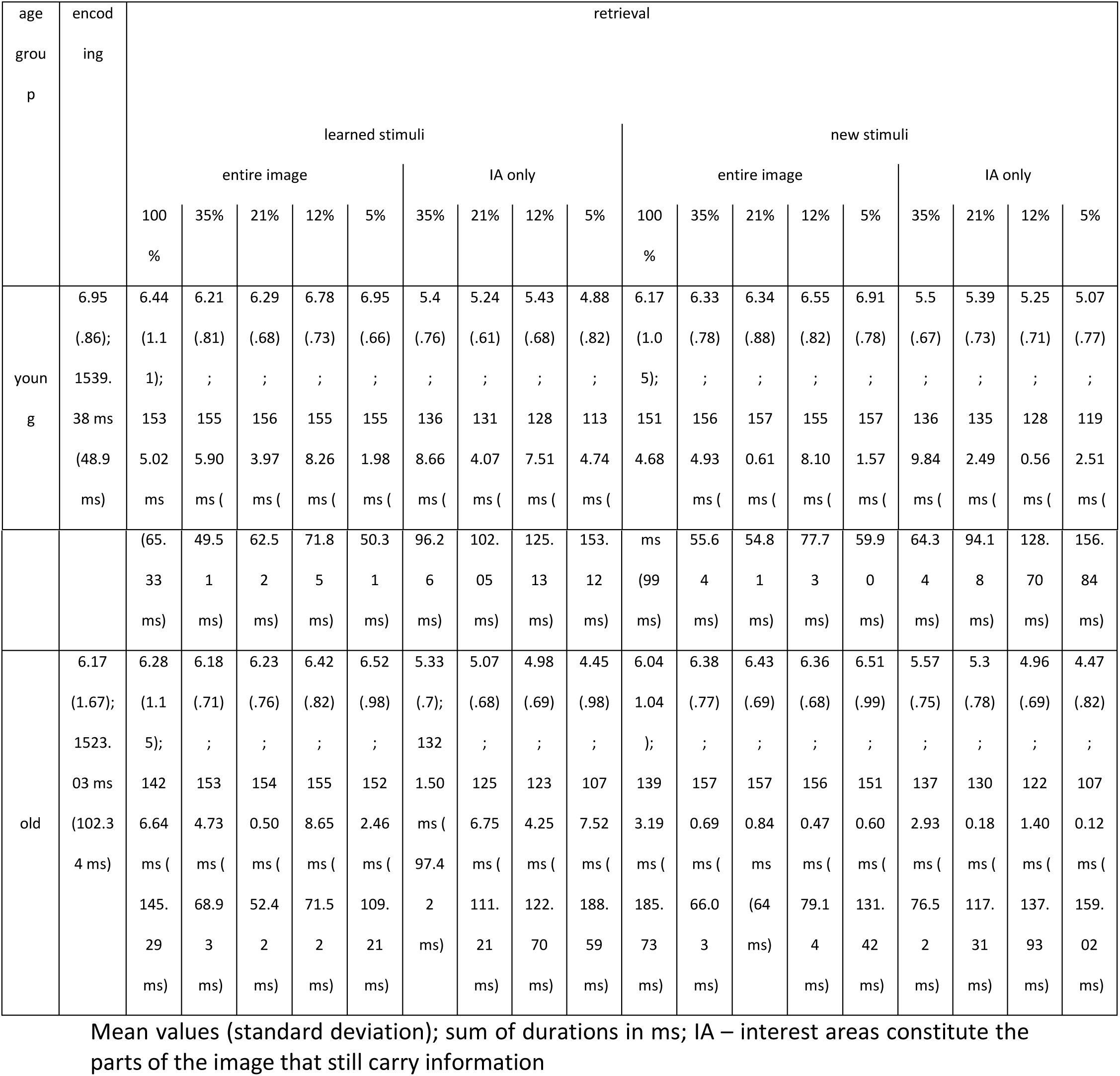
Experiment 1 - Fixation numbers and durations during encoding, and during retrieval per stimulus type and completeness level and within specific interest areas.

#### Fixations during retrieval

First, we inspected the overall number of fixations in a similar way as for behavioral accuracy. We subjected the fixation numbers to a three-way mixed ANOVA with factors age (young, old), stimulus completeness (100%, 35%, 21%, 12%, 5%) and stimulus type (learned, new). Interestingly, young and older adults did not differ in how much they fixated on the images (see Table 3; main effect of age: *F*_(1,48)_ = 0.604, *p* = .441, *BF_01_* = 2.184), nor was there a difference between learned and new stimuli (main effect of stimulus type: *F*_(1,48)_ = 0.480, *p* = .492, *BF_01_* = 8.150). However, participants fixated slightly more when less of the image was visible (main effect of stimulus completeness: *F*_(4,192)_ = 11.083, *p* < .001, *BF_01_* = 0.086).

Secondly, there was an inverse relationship between the number of fixations and their average duration, i.e., the longer participants spent fixating on one location, the lower the number of fixations was and vice versa. Given that participants only had two seconds to look at each image, this reflects a trade-off between how many locations to look at and how long to look at each of them. Therefore, we analyzed the sum of durations for all fixations rather than average fixation duration. While there were no main effects for age or stimulus type (main effect of age: *F*_(1,48)_ = 3.189, *p* = .08, *BF_01_* = 0.823; main effect of stimulus type: *F*_(1,48)_ = 0.442, *p* = .510, *BF_01_* = 8.855), stimulus completeness affected fixation durations (main effect of stimulus completeness: *F*_(4,192)_ = 24.035, *p* < .001, *BF_01_* = 1.170e-20), and interacted with age (age × stimulus completeness: *F*_(4,192)_ = 8.671, *p* < .001). More specifically, older adults fixated only shortly on the full stimuli but spend more time fixating the masked stimuli with durations similar to those of young adults whose fixation durations did not differ across masking levels.

#### Fixations in IAs

To identify areas of the images at which participants were looking, we examined fixations in predefined IAs. As explained in the methods, the stimuli were masked at different levels, so that only parts of the image were visible. We tested whether the age groups showed differential fixation patterns at the relevant positions, i.e., the parts of the image that still carried information, which might explain the observed age-related response bias. Thus, IAs comprised all the unmasked portions of the image with an added margin of 25 pixels (to include areas in the foveal focus).

Again, as revealed by a three-way mixed ANOVA, young and older adults did not differ overall in their number of fixations on IAs (see Table 3; main effect of age: *F*_(1,48)_ = 1.497, *p* = .227, *BF_01_* = 2.092), nor were there differences between learned and new stimuli (main effect of stimulus type: *F*_(1,48)_ = 0.288, *p* = .594, *BF_01_* = 9.522). Both groups had less fixations on less complete images (main effect of stimulus completeness: *F*_(4,192)_ = 66.497, *p* < .001, *BF_01_* = 3.513e-59). That was to be expected, because the IAs got smaller with increasing masking levels. The same was true for fixation durations, i.e., all participants fixated shorter on smaller IAs and spent more time fixating on the more complete stimuli (main effect of stimulus completeness: *F*_(4,192)_ = 125.125, *p* < .001, *BF_01_* = 7.017e-88), and there were no differences between learned and new stimuli (main effect of stimulus type: *F*_(1,48)_ = 2.223, *p* = .143, *BF_01_* = 7.427). However, here younger adults fixated slightly longer on the images than older adults (main effect of age: *F*_(1,48)_ = 6.826, *p* = .012, *BF_01_* = 0.343).

#### Spatial distribution of fixations

In addition to investigating the number and duration of fixations, we also tested whether different viewing patterns could drive recognition performance.

Given that the new stimuli in particular were often erroneously recognized as one of the learned stimuli by older adults, we computed the spatial overlap of fixations between conditions. Specifically, we calculated two-dimensional correlation coefficients between fixation heatmaps (see methods eye-tracking analysis) of correctly identified learned stimuli (e.g. library identified as library) with falsely identified stimulus-matching stimuli (e.g. library identified as kitchen), and with falsely identified response-matching stimuli (e.g. office identified as library), separately for all levels of stimulus completeness (for exemplary heatmaps and how the correlations were calculated, see Figure 1D). The resulting *r*-values were Fisher’s *z*-transformed and averaged across stimulus identity and completeness, resulting in one response-matching and one stimulus-matching correlation value per participant. These values were subjected to a two-way mixed ANOVA with factors age group and match type. There were no differences between age groups (main effect of age: *F*_(1,48)_ = 0.002, *p* = .965, *BF_01_* = 4.166), but correlations for stimulus-matching heatmaps were considerably higher than for response-matching heatmaps (main effect of match type: *F*_(1,48)_ = 194.381, *p* < .001, *BF_01_* = 5.074e-21), suggesting that the viewing patterns were driven by the stimulus rather than by the response choice. This effect also interacted with age (age × match type: *F*_(1,48)_ = 9.332, *p* = .004). Post-hoc t-tests revealed that older adults’ fixation patterns for response-matching stimuli correlated more than those of young adults (*t*_(48)_ = − 2.882, *p* = .006, *BF_01_* = 0.136), but there were no group differences between stimulus-matching heatmap correlations (*t*_(48)_ = 1.512, *p* = .137, *BF_01_* = 1.397). To test the heatmaps’ relation to the suggested pattern completion bias, we correlated the heatmap match scores with the cumulative partial response bias (Pearson correlation). Both the stimulus-matching and the response-matching score positively correlated with the response bias in older adults (stimulus-matching: *r* = 0.483, *p* = .017; response-matching: *r* = 0.418, *p* = .042), but not in young adults (stimulus-matching: *r* = 0.005, *p* = .981; response-matching: *r* = 0.346, *p* = .084), indicating that the overlap of fixation heatmaps was related to the age-related pattern completion bias.

#### Summary of the eye-tracking results

During encoding, older adults had slightly lower fixation numbers than young adults, however, this difference was not predictive of the observed age-related response bias. Both age groups had similar fixation patterns during retrieval albeit some differences in fixation durations. Specifically, older adults had shorter viewing durations than young adults for complete stimuli and within IAs. Given the appearance of a masked stimulus, one may argue that the observed viewing pattern was a useful strategy for this task. That is, with decreased stimulus completeness there was less to see at each fixation, rendering it necessary to shift fixations more often to obtain a similar amount of information as for the unmasked stimuli. Importantly, there were no age-related viewing differences between learned and new stimuli which could account for the observed response bias. Analyses of fixation heatmaps revealed that viewing patterns were more stimulus-driven than predictive of response choices. That is, independent of recognition, fixation patterns for e.g. ‘library’ trials were more similar to other ‘library’ trials than fixation patterns for ‘office’ trials that were falsely recognized as ‘library’. Interestingly though, within the response-matching heatmap comparison, older adults had higher values than young adults, which also correlated with the response bias. These results suggest that the spatial overlap between fixations was related to the response bias in older adults, i.e., the more fixation patterns overlap between correctly and incorrectly identified stimuli, the stronger the bias to choose a familiar response and to falsely recognize a new stimulus.

## Experiment 2

### Methods

To obtain longitudinal performance profiles and detect potential performance changes in clinical studies, it is necessary to test participants on multiple occasions. We therefore aimed at developing parallel versions of the MIC, comparable in difficulty. We used fewer stimuli to facilitate quick administration often necessary in clinical studies. In addition, a subset of participants was tested twice on different versions of the task to assess the reliability of the task and to investigate potential learning effects independent of the stimuli.

#### Subjects

All participants were recruited from existing databases at the DZNE in Magdeburg. Sixty young and healthy adults (19-35 years; 43 females) participated in experiment 2, and were randomly assigned to one of four groups pertaining to a specific version of the task (15 participants per task version). Thirty-six of the participants came in for a second time and were again distributed across versions (nine participants per task version). Informed written consent was obtained before the experiment, and the study received approval from the Ethics Committee of the University of Magdeburg. All participants received monetary compensation of 6.50€/h.

#### Materials

Forty-eight black and white line-drawn images of simple indoor scenes were created using Autodesk 3DS Max. Four different sets of 12 stimuli each (4 to be learned, 4 to test learning, 4 new) constituted different versions of the task (see exemplary stimuli for two sets in Figure 2A). Again, the stimuli were gradually masked (stimulus completeness: 100%, 35%, 21%, 12%, 5%). In total, there were 40 stimuli in the test phase of each version, and each stimulus was repeated four times.

**Figure 2.**
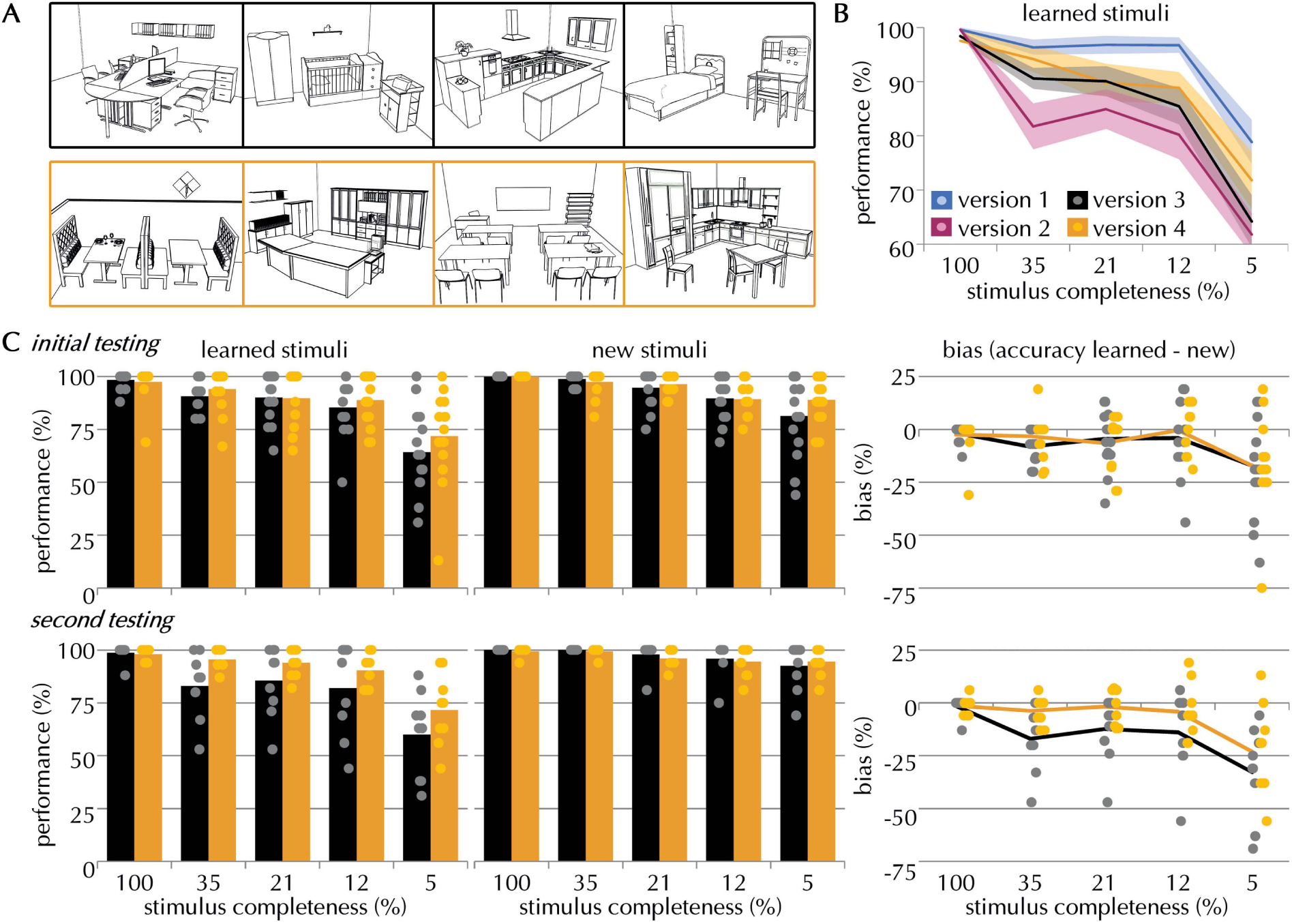
- Experiment 2. **(A)** Exemplary learned stimuli for versions 3 (black) and 4 (orange). **(B)** Performance for learned stimuli separately for all tested versions (blue: version 1, magenta: version 2, black: version 3, orange: version 4) and the 5 different levels of stimulus completeness (mean ± standard error). **(C)** Performance and bias measures at initial (top) and second testing (bottom) for comparable versions 3 and 4; left: performance separately for learned and new stimuli for the 5 different levels of stimulus completeness (mean); right, bias measure (see methods for a detailed explanation) - difference in accuracy scores for learned minus new stimuli calculated separately for each participant (mean); positive values indicate a bias toward pattern completion.

#### Procedure

We administered the MIC as described in experiment 1 but in four different versions. Apart from containing a different set of stimuli, all task versions were identical. Fifteen participants each were randomly assigned to one of the four versions. In each version, participants had to learn four different scene exemplars, resulting in a chance level of 1/5 in the test phase. Thirty-six of the participants performed two versions of the task with a minimum of two weeks in-between tests. All possible pairings of tasks were tested in a between-subject design (e.g. versions 1-2, 1-3, 1-4, 2-1, 2-3, etc.).

#### Exploration of task differences using Bayesian analyses

As in experiment 1, we report Bayes factor *BF_01_* in favor of the null hypothesis to estimate support for the similarity of the task versions.

## Results

### Results at initial testing

In order to compare the four different versions, performance data of all initial tests (N = 60) were submitted to a three-way mixed ANOVA with between-subjects factor version (1-4), and within-subjects factors stimulus completeness (100%, 35%, 21%, 12%, 5%) and stimulus type (learned, new). Performance across all four versions differed significantly (see Figure 2B; main effect of version: *F*_(3,56)_ = 3.935, *p* = .013). Reduced stimulus completeness resulted in less accurate performance (main effect of stimulus completeness: *F*_(4,224)_ = 102.147, *p* < .001), and performance was better for new than for learned stimuli (main effect of stimulus type: *F*_(1,56)_ = 35.826, *p* < .001). Additionally, there was an interaction between version and stimulus completeness but no other interactions were significant (version × stimulus completeness: *F*_(12,224)_ = 2.616, *p* = .003; version × stimulus type × stimulus completeness: *F*_(12,224)_ = .911, *p* = .537, version × stimulus type: *F*_(3,56)_ = 1.949, *p* = .123). Importantly though, as analyzed by a separate ANOVA, response biases did not differ between versions (main effect of version: *F*_(3,56)_ = 1.949, *p* = .132).

Descriptive data (see Figure 2B) showed that versions 3 and 4 yielded similar results, whereas version 1 seemed to be easier and version 2 seemed to be more difficult. As we wanted to obtain comparable versions, we continued analyses only with data from versions 3 and 4 (N = 30). Indeed, there was no overall difference between these two versions (see Figure 2C top panel; main effect of version: *F*_(1,28)_ = 0.781, *p* = .384, *BF_01_* = 3.586), nor were there any interactions with the other factors (version × stimulus type: *F*_(1,28)_ = 0.098, *p* = .757; version × stimulus completeness: *F*_(4,112)_ = 1.776, *p* = .139; version × stimulus type × stimulus completeness: *F*_(4,112)_ = 0.569, *p* = .685). Other than that, participants were less accurate with decreasing stimulus completeness (main effect of stimulus completeness: *F*_(4,112)_ = 54.995, *p* < .001, *BF_01_* = 4.351e-25) and performed better for new than for learned stimuli (main effect of stimulus type: *F*_(1,28)_ = 12.918, *p* = .001, *BF_01_* = 7.407e-4). Furthermore, the response bias did also not differ between versions (main effect of version: *F*_(1,28)_ = 0.098, *p* = .757, *BF_01_* = 3.242; version × stimulus completeness: *F*_(4,112)_ = 0.569, *p* = .685), but it slightly decreased with decreasing stimulus completeness (main effect of stimulus completeness: *F*_(4,112)_ = 9.646, *p* < .001, *BF_01_* = 4.013e-5). This was confirmed by one-sample t-tests against 0, revealing that there was a negative response bias at 5% stimulus completeness, but at no other completeness level (after Holm-Bonferroni multiple comparisons correction, level 100%: *t*_(14_) = −1.718, *p* = .108; level 35%: *t*_(14)_ = −1.624, *p* = .127, level 21%: *t*_(14)_ = −1.695, *p* = .112; level 12%: *t*_(14)_ = −0.569, *p* = .578; level 5%: *t*_(14)_ = −3.439, *p* = .004).

### Results at second testing

Performance data of all participants who came in for a second test of versions 3 and 4 (N = 18) were submitted to a mixed ANOVA with the factors task version (3, 4), stimulus completeness (100%, 35%, 21%, 12%, 5%) and stimulus type (learned, new). Most importantly, the versions did not differ from each other (see Figure 2C bottom panel; main effect of version: *F*_(1,16)_ = 1.634, *p* = .219, *BF_01_* = 2.306), nor were there any interactions (version × stimulus type: *F*_(1,16)_ = 3.001, *p* = .102; version × stimulus completeness: *F*_(4,64)_ = 1.779, *p* = .144; version × stimulus type × stimulus completeness: *F*_(4,64)_ = 0.961, *p* = .435). Reduced stimulus completeness resulted in less accurate performance (main effect of stimulus completeness: *F*_(4,64)_ = 42.952, *p* < .001, *BF_01_* = 4.742e-8), and performance was better for new than for learned stimuli (main effect of stimulus type: *F*_(1,16)_ = 19.776, *p* <.001, *BF_01_* = 3.921e-7). Additionally, response biases did not differ between versions (main effect of version: *F*_(1,16)_ = 3.001, *p* = .102, *BF_01_* = 1.083; version × stimulus completeness: *F*_(4,64)_ = 0.961, *p* = .435), but they were affected by stimulus completeness (main effect of stimulus completeness: *F*_(4,64)_ = 14.793, *p* < .001, *BF_01_* = 6.334e-7). Arguably though, *BF_01_* for the factor version was around 1, indicating that no strong conclusion can be drawn about the version-specific difference of the response biases during second testing; descriptively, version 3 may show a slight negative bias in relation to version 4 (i.e. a bias toward pattern separation; see Figure 2C), nevertheless, this is not backed up statistically.

### Comparison between time points

To be able to measure change over time, it is important that the parallel versions of the MIC are not sensitive to general practice or learning effects. Due to the between-subjects design across all four versions, we only had six participants who completed version 3 or 4 at first and second testing However, given that general learning effects should be independent of the particular version applied at the first testing session, we realized the time point analysis in a between-subjects design. Specifically, we compared performance between participants who were tested with versions 3 or 4 in the initial session (N = 15) with a different set of participants who were tested on versions 3 or 4 in the second session (N = 15). The six participants who had been tested on both versions were equally distributed over these two groups (three in each group; total per group N=18). To keep group size equal, only participants who came in twice were included here.

A mixed ANOVA with the between-subjects factor time (initial or second test), and the within-subjects factors stimulus type and completeness revealed no overall learning effects between time points (see Table 4; main effect of time: *F*_(1,28)_ = 1.408, *p* = .245, *BF_01_* = 2.970), nor were there any interactions (time × stimulus type: *F*_(1,28)_ = 0.389, *p* = .583; time × stimulus completeness: *F*_(4,112)_ = 2.068, *p* = .090; time × stimulus type × stimulus completeness: *F*_(4,112)_ = 0.637, *p* = .637). Effects of stimulus type and completeness followed the same profile as before (main effect of stimulus completeness: *F*_(4,112)_ = 75.683, *p* < .001, *BF_01_* = 2.748e-23; main effect of stimulus type: *F*_(1,28)_ = 21.807, *p* < .001, *BF_01_* = 1.279e-6). Importantly, response biases also did not change over time (main effect of time: *F*_(1,28)_ = 0.389, *p* = .538, *BF_01_* = 3.090; time × stimulus completeness: *F*_(4,112)_ = 0.637, *p* = .637), but were again affected by stimulus completeness (main effect of stimulus completeness: *F*_(4,112)_ = 17.317, *p* < .001, *BF_01_* = 2.214e-9).

**Table 4.**
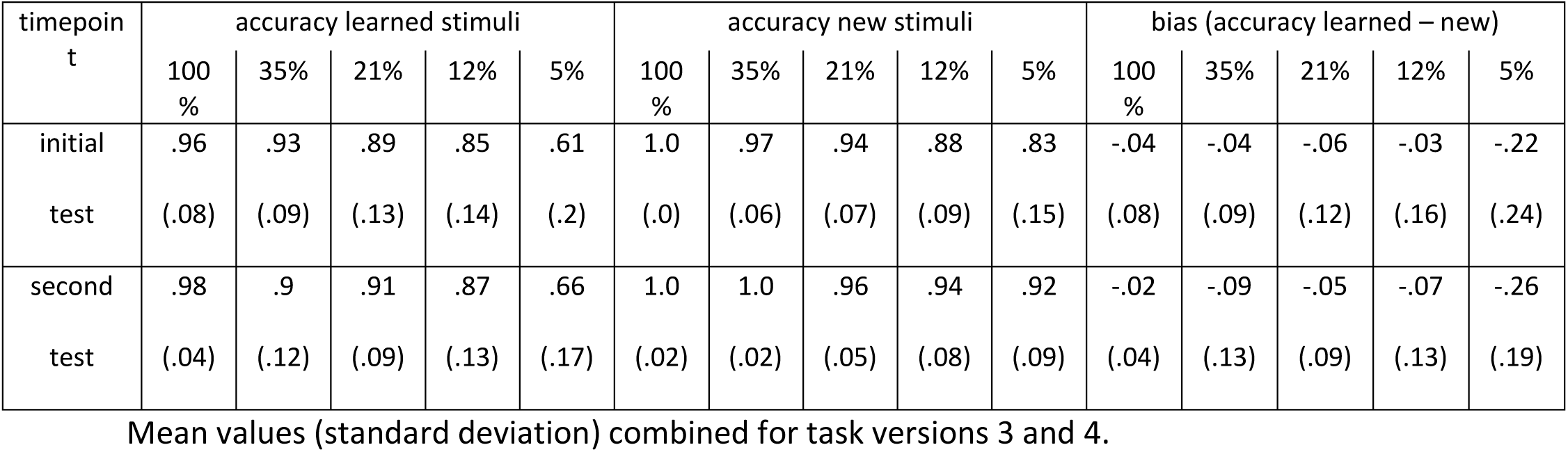
Experiment 2 - Performance results for multiple timepoint testing per stimulus type and completeness levels.

### Summary of the results

Out of the four developed stimulus sets, versions 3 and 4 yielded comparable results which followed the same performance profiles as the original MIC, independent of whether they were tested for the first or a second time. We did not observe any learning effects when participants were tested twice on the task. Note that there were no differences in reaction times between versions or time points either (see Supplementary material).

## Discussion

In this study, we could replicate the behavioral findings of our earlier paper (Vieweg et al., 2015). Specifically, recognition memory declined with reduced stimulus completeness, more so in aging, and older adults chose familiar responses over new ones, suggesting a bias towards pattern completion. Associated eye-tracking data of older adults showed fewer fixations during encoding and slightly shorter IA fixation durations during retrieval but with no distinction between learned and new stimuli. While this particular difference may have contributed to the overall performance reduction of older adults, it could not account for the more specific impairment in the recognition of new items, i.e. although the age groups showed small encoding differences, the age-related pattern completion bias could not be attributed to this difference. However, when a new stimulus was falsely identified as a learned stimulus, the corresponding fixation patterns overlapped more in older adults than in young adults, and only in older adults did the resulting score correlate with the performance bias suggesting a relation between the two. Nevertheless, spatial viewing patterns for both age groups were more driven by stimulus identity and not so much by response choice. Thus, the observed differential recognition memory effects cannot solely be explained by the corresponding eye-movements, lending support for our hypothesis that differences in pattern completion are accountable for the observed effects.

In a second experiment, we developed four distinct and shorter versions of the MIC to provide a means for repeated testing. Given the complexity of scene images, it is very likely that they differ in their recognizability, which is further influenced by the set within which they are situated. For example, the recognizability of the scene ‘kitchen’ does not only depend on the image itself, but also on the visual similarity of other images from the same set of stimuli. Thus, it is far from surprising that different task versions differ in difficulty. Nevertheless, of the four tested versions, two tasks proved to be of similar difficulty, and performance for these two tasks was comparable across two time points, suggesting that there was no general learning effect of the task that could distort performance profiles. Consequently, we can now provide two task versions of comparable difficulty for repeated testing.

The previous literature on memory-related viewing behavior has linked increased fixation numbers during encoding to better memory retrieval (Loftus, 1972). With respect to that, older adults’ fewer fixation numbers during encoding may be similarly related to their impaired retrieval – an issue that was also raised by Molitor and colleagues (2014) who found that false alarm trials were associated with worse encoding fixation patterns than correctly identified stimuli. However, here, correlations of eye-tracking data during encoding with pattern completion ability or bias were not observed. Consequently, performance differences could not be predicted from differential eye movements during encoding, suggesting the involvement of another process, which we suggest to be an age-related pattern completion bias.

Further on, fixation durations at informative positions (i.e. IAs) during retrieval were slightly longer in the younger age group, potentially accounting for their overall recognition advantage. This idea receives support from previous observations that longer fixations during retrieval would code for a prior exposure indicating recognition (Hannula et al., 2012; Molitor et al., 2014). However, reports in the literature also show that saccade velocity and reaction times (Moschner and Baloh, 1994) as well as saccade frequencies and amplitude (Dowiasch et al., 2015) decrease with age, in line with a general slowing of processing speed (O’Shea et al., 2016). Based on these findings, shorter fixation durations in older adults could be explained by their increased saccade durations, i.e. eye-movements between fixations. Thus, shorter fixation durations and lower fixation numbers might well account for recognition memory deficits, however, they do not provide a reliable index of the specificity of the underlying process, be it processing speed or memory decline. Nevertheless, to prevent potential encoding differences related to processing speed in the future, we have adjusted our paradigm in such a way that during the learning phase participants can decide in each trial how long the stimulus is presented.

In further support of recognition-rather than perceptual differences, the main age-related disparity in this task was observed in the difference in performance for learned and new stimuli, which was absent in the eye-tracking data. Hence, while older adults made more errors for new stimuli than for learned ones, their eye-movements were not differentiable between the two stimulus types. Nevertheless, the age group interaction for spatial overlap of fixations is noteworthy; although fixation patterns were generally more stimulus-than response-driven, older adults’ fixations for new stimuli - which were identified as old - shared more overlap with the corresponding old stimuli than was the case for young adults. This observation partly supports what we were assuming from the false alarm distributions, namely that for incorrect recognition, participants chose the stimulus they perceived as the most similar. However, given the distinctly fewer errors in young adults, their heatmap correlations consist of considerably fewer trials, which inevitably affects their overall correlation score, and consequently puts some limits to the interpretability of these scores. In summary, the specificity of the older adults’ errors, i.e. more often picking a specific familiar response instead of the ‘none of these’ option in identifying new stimuli, and the absence of a clear corresponding viewing pattern, lends further credibility to the interpretation that age differences observed with this task are a result of a pattern completion bias.

It is also worth noting that the behavioral findings could almost completely be replicated from their first implementation (Vieweg et al., 2015). The only deviation was a non-significant group difference in the bias score for the 12% completeness level, but the overall shape of the curve was the same, and so were all other results concerning accuracies, false alarm distributions, reaction times, and confidence ratings (see Supplementary for the latter two). This provides compelling evidence that the task in its current form can be reliably applied and the obtained results are not coincidental. Thus, the MIC can be used to pick up fine-grained age-related recognition memory differences in the scheme of pattern completion, which are in line with models of cognitive aging (Wilson et al., 2006). The MIC provides two important measures: pattern completion ability – quantified by recognition performance for learned stimuli, and a pattern completion bias – quantified by the interplay between learned and new images. Furthermore, we have tested and can now provide additional, parallel versions of the MIC whose performance profiles match the original task, but which are fast and easy to administer. The MIC, including all parallel versions, is now publicly available on the open science framework (https://osf.io/juvwy) in order to advance the research field and to enable other scientists to employ a pattern-completion-specific task in their studies. We believe that the MIC could prove particularly useful for longitudinal studies (i.e. on lifespan development and dementia), or as a clinical tool in patient and intervention studies.

## Acknowledgments

Parts of this manuscript have been used in different form in P.V.’s doctoral thesis (Vieweg, 2017). P.V. and T.W. are supported by European Research Council Starting Investigator Grant AGESPACE (335090). M.R. is supported by European Social Fonds (ESF) (Sachsen-Anhalt Wissenschaft Spitzenforschung/Synergien: AGETIME).

We would like to thank Henrike Raith, Franziska Schulze, Anica Luther, Patrick Hauff and Sophie Schnau for their help with data collection and neuropsychological testing, and Selim Candan for help with stimulus design.

